# Disruption of immunoglobulin heavy and light chain assembly by antisense oligonucleotides impairs protein homeostasis and myeloma cell survival

**DOI:** 10.1101/2023.08.29.555385

**Authors:** Catherine Horiot, Murielle Roussel, Antoine Praité, Justine Pollet, Sébastien Bender, Christophe Sirac, Amélie Bonaud, Laurent Delpy

## Abstract

Multiple myeloma (MM) is related to the accumulation of malignant plasma cells (PCs) in the bone marrow. MM accounts for approximatively 10% of hematological malignancies and despite major improvement in therapies and outcomes, relapses will virtually occur in all patients. Usually, the disease goes along with an excess production of a monoclonal immunoglobulin (Ig) component by the tumor PC clone. However, many questions remain regarding the consequences of a deregulated Ig production on PC survival. Recent advances in RNA-based therapy using antisense oligonucleotides (ASO) prompted us to examine the impact of altered Ig heavy to light chain (HC/LC) ratios in MM cells. We designed a pan IgG subclasses specific ASO targeting a consensus sequence found in the polyadenylation signal (PAS) of all secreted *IGHG* mRNAs (IgG-ASO). Remarkably, treatment with this compound strongly decreased IgG secretion in MM cell lines and patient cells. Consistent with a deregulated HC/LC ratio, a dose-dependent excess of free-LCs (as monomers and dimers) was observed in myeloma cells treated with IgG-ASO, compared to an irrevelant control ASO (CTRL). RNA-seq profiles further indicated that the expression of genes involved in cellular metabolism, unfolded protein response (UPR) and cell death pathways were altered after treatment with IgG-ASO. Interestingly, impaired survival of primary IgG-expressing cells isolated from MM patients was achieved upon treatment with IgG-ASO, whereas no major effect was observed for healthy cells. Altogether, our data provide evidence for efficient inhibition of IgG secretion upon ASO treatment and suggest that an excess of free-LC due to disruption of HC/LC stoichiometry is toxic for MM cells expressing complete Ig. Such RNA-based strategies targeting PC in an Ig isotype-dependent manner could open new avenues for selective therapeutic approaches in PC dyscrasias.

## INTRODUCTION

Multiple Myeloma (MM) is a plasma cell (PC) neoplasm characterized by the presence of tumor clones in bone marrow niches. The abnormal proliferation of PC clones is accompanied by excess production of a monoclonal immunoglobulin (Ig), as serum M-component, in most of the cases. MM accounts for approximatively 10% of all hematological malignancies and the risk of developing this disease increases with age, with a median age at diagnosis > 65 years (Kumar et al., 2014; Moreau, 2017). In the majority of patients, pre-tumoral and/or asymptomatic stage called MGUS (Monoclonal Gammapathy of Undetermined Significance) and smoldering MM (SMM), respectively, can be detected years before the diagnosis of symptomatic MM. The risk of progression from MGUS to MM is around 1% per year (Rajkumar, 2022). Although continued improvement in the survival of myeloma patients has been observed over the last decades (Kumar et al., 2014), MM remains uncurable with relapses eventually occurring in nearly all patients. Alternative therapeutic approaches are still needed (van de Donk et al., 2021). Currently, new immunotherapies using antibodies targeting molecules strongly expressed by tumor cells, as CD38 (Daratumumab, Isatuximab), BCMA (Belantamab Mafodotin), or SLAMF7 (Elotuzumab) have significantly improve MM patients outcome. The transmembrane glycoprotein BCMA is also the target of CAR-T cell therapy for relapsed/refractory patients with remarkable efficacy (Van Oekelen et al., 2023).

PCs are real Ig production factories capable of secreting up to 10000 Ig/sec (Cenci et al., 2011; Eyer et al., 2017). These antibody-secreting cells owe their survival to a very tight control of proteostasis through activation of the unfolded protein response (UPR), a signaling pathway that adjusts the capacity of the endoplasmic reticulum (ER) and promotes protein folding, while increasing the degradation of unfolded/misfolded proteins by the proteasome. Accordingly, modulation of intracellular proteostasis is an attractive therapeutic approach and proteasome inhibitors (PIs) was introduced in the treatment landscape of MM patients and those suffering from monoclonal gammopathies with clinical significance (MGCS) over the last decades (Gandolfi et al., 2017; Jaccard et al., 2014; Cohen et al., 2015; Fermand et al., 2018). In addition, apoptosis induced upon proteasome inhibition was correlated with the amount of Ig production (Meister et al., 2007). Consistent with a major role for Ig production in myeloma cell survival, a positive correlation between the amount of intracellular Ig light chains (LC) in tumor PCs and progression free survival (PFS) has been recently observed in newly diagnosed MM patients (Vdovin et al., 2022).

IgG (60%) and to a lesser extent IgA (24%) were the most frequent paraproteins expressed by myeloma cells, in a large cohort study of 10,750 patients. A minority of PC clones expressed only LC (11%) or IgD (3%) (Greipp et al., 2005). Usually, PCs make more LC than heavy chain (HC) (Katzmann et al., 2002), and excessive production of free-LC is a characteristic feature of PC dyscrasias (Sirac et al., 2021). The deregulation of Ig production has emerged as a potential new targeted therapy for MM and other PC disorders. By using siRNA knockdown strategies, Zhou *et al* have shown that the reduction of LC but not HC impaired the proliferation of tumor PC-expressing complete Ig (Zhou et al., 2014). However, many questions remains regarding the impact of disrupted HC/LC ratios in PC survival.

Antisense oligonucleotides (ASOs) are short single strand of modified DNA that can be used as “gapmer ASOs” to degrade RNA by RNAse H1 recruitment, or as “steric blocker ASOs” to modify RNA maturation processes such as splicing or polyadenylation, depending of the oligonucleotide chemistry (Bennett, 2019). In the lab, we previously used steric blocker ASOs to modify Ig expression in B-lineage cells (Ashi et al., 2018; Marchalot et al., 2020, 2021). Here, we assessed the impact of an ASO-mediated decrease in HC expression in IgG-expressing myeloma cells. We provide evidence for efficient inhibition of IgG secretion, and that disruption of HC/LC stoichiometry impairs the survival of myeloma cells due to altered protein homeostasis.

## MATERIAL & METHODS

### ASO design

The *IGH* gene sequence was collected from NCBI (GRCh38.p12 assembly) and secreted polyadenylation signal (PAS) sequences from all HC subclasses were aligned using Multalin version 5.4.1 software (Fig. S1) (Corpet, 1988). We designed an ASO targeting a consensus RNA sequence present in all secreted *IGHG* PAS, as a pan-*IGHG* subclasses ASO (IgG-ASO or referred as ASO for figures: 5’-GCGCTGGGTGCTTTATTTCCATG-3’). Both IgG-ASO and an irrelevant control (CTR: 5′-CCTCTTACCTCAGTTACAATTTATA-3′) were designed as Vivo-morpholinos (*i.e.* morpholino oligos coupled to an octa-guanidine dendrimer by their 3’ part) and purchased at Gene Tools, LLC. Stock solutions were resuspended at 0.5mM in sterile nuclease-free water and stored at room temperature protected from light.

### Cell culture and ASO treatments

One IL-6-independent (LP1) and two IL-6-dependent (XG2 and XG6) myeloma cell lines expressing a monoclonal IgG *Lambda* (λ) isotype were used. LP1 cells were cultured in Iscove medium (IMDM) with stable Glutamine and 25 mM HEPES (Biowest) and supplemented with 20% Fetal Bovine Serum (FBS) (Dominique Dutscher). For XG2 and XG6 cells, recombinant human IL-6 (2 ng/mL) (Proteintech) was added to RPMI1640 culture medium containing Ultraglutamine (Lonza) and 10% FBS. MM patient’s cells collected from bone marrow aspirates after a Ficoll density gradient separation (Lympholyte®-H Cell Separation Media, Cedarlane). Primary cells were cultured in RPMI1640 supplemented with 10% FBS, Penicillin/Streptomycin and IL-6 (2 ng/mL). For ASO treatments, cells were seeded at 5×10^5^ to 1×10^6^ cells/ml and ASO were added (1 to 6 µM) for 48h or as otherwise indicated. All cell culture was performed at 37°C in a humidified 5% CO2 atmosphere.

### Cell viability

Cell viability was usually monitored after trypan blue coloration. MTS [3-(4,5-dimethylthiazol-2-yl)-5-(3-carboxymethoxyphenyl)-2-(4-sulfophenyl)-2H-tetrazolium, inner salt] cell proliferation and viability assays (CellTiter 96® AQueous One Solution Cell Proliferation Assay, Invitrogen) were performed according to the manufacturer’s instructions with 50 000 cells per well and read at 492nm on a microplate absorbance reader (Multiskan FC microplate photometer, Thermo Fisher).

### Flow cytometry

For FACS analysis on primary cells treated or not with ASO for 48h, cell suspensions were washed in phosphate-buffered saline (PBS) and labelled with mouse anti-human CD38 and CD138 mAbs (Table S1) for 30 minutes in FACS buffer (PBS supplemented with 2% FBS and 2 mM EDTA). To exclude non-viable cells, 0.2 µl of FVS (Fixable Viability Stain, BD Horizon, ref 564406) was added during labelling. Data were acquired on a CytoFLEX flow cytometer (Beckman Coulter Life Science) and analysed using Flowlogic^TM^ software (Miltenyi Biotec).

### Western blot

Cells were lysed in radioimmunoprecipitation assay (RIPA) buffer (Thermo Scientific) containing a cocktail of protease and phosphatase inhibitors. Proteins were quantified with Pierce™ BCA Protein Assay kit (Thermo Scientific). After a denaturation at 95°C during 6 minutes, samples were separated on 12% SDS-PAGE TGX Stain-Free FastCast Acrylamide gels (Bio-Rad Laboratories). Proteins were then electro-transferred onto PVDF (polyvinylidene fluoride) membranes with Trans-Blot Turbo transfer system (Bio-Rad Laboratories). Antibodies used for the detection are listed in Table S1. Detection of HRP-linked antibodies was performed using chemiluminescence detection kits (ECL™ or ECL Plus™, GE Healthcare) according to the manufacturer’s instructions and visualized using ChemiDoc™ Touch Imaging System (Bio-Rad Laboratories). Image Lab Software (Bio-Rad Laboratories) was used to quantify the bands.

### RNA extraction and RT-qPCR

Total RNA was prepared using Trizol and RNA Direct-ZOL procedures (Zymo Research R2062). Reverse transcription (RT) was performed on 200 ng to 1 µg DNase I-treated RNA using High-Capacity cDNA Reverse Transcription Kit (Applied Biosystems). Quantitative PCRs (qPCRs) were performed on cDNA samples using SensiFAST™ SYBR® Hi-ROX or SensiFAST™ Probe No-ROX kits (Bioline) on a StepOnePlus real-time PCR apparatus (Applied Biosystems). Transcripts were quantified according to the standard 2^-ΔΔCt^ method after normalization to GAPDH (Hs.PT.39a.22214836, Integrated DNA Technologies). Primers used for SYBR® technology were manufactured by Eurofins Genomics and listed in Table S2.

### Enzyme-linked immunosorbent assay (ELISA)

Human IgG and Igλ concentrations were determined in culture supernatants and cell pellets as indicated. Sandwich ELISAs were performed in polycarbonate 96-multiwell plates using polyclonal goat anti-human IgG or Igλ antibodies (Southern Biotechnologies) (Table S1). Alkaline Phosphatase (AP) activity was assayed using SIGMAFAST^TM^ nitrophenyl phosphate tablets (Sigma-Aldrich) and the reaction was blocked with the addition of NaOH. Optic density was then measured at 405 nm on a Multiskan FC microplate photometer (Thermo Scientific).

### Quantification of Ig Lambda Free-light chains

Quantification of Lambda Free-Light Chains was performed in culture supernatants using the Optilite® Freelite® Lambda Free kit (The Binding Site™) in a Optilite turbidimeter (The Binding Site™).

### RNA preparation and sequencing

Total RNA was prepared from XG6 cells treated or not with IgG-ASO or CTR (48h, 3µM) using the NucleoSpin® RNA L kit (Macherey-Nagel, France) according to manufacturer instructions. Samples quality controls and libraries preparation were performed at the GeT-Santé facility (Inserm, Toulouse, France, get.genotoul.fr). Libraries quantification and sequencing were then performed at the GeT-PlaGe core facility (INRAE, Toulouse, France), as described (Thomas et al., 2023). Briefly, libraries were equimolarly pooled and RNA sequencing was performed on one S4 lane of the Illumina NovaSeq 6000 instrument (Illumina, San Diego, USA), using the NovaSeq 6000 S4 v1.5 Reagent Kit (300 cycles), and a paired-end 2 x 150 pb strategy. A total of 1.5 billion reads was obtained.

### Bioinformatic analyses

Paired-end reads were processed through the bioinformatic pipeline nf-core/rnaseq v3.12 (Patel et al., 2023) in a reverse strandedness mode. Quality control and trimming were performed via fastp (Chen et al., 2018), upon alignment on GRCh38 human genome with STAR (Dobin et al., 2013) and quantification by Salmon (Patro et al., 2017). The resulting matrix of gene-level raw counts were used with DESeq2 (Love et al., 2014) to extract differentially expressed (DE) genes (adjusted *p* value < 0.01 and absolute log2foldchange > 0.5) for comparison and downstream analysis. Volcano plot were drawn with EnhancedVolcano R package (Blighe et al., 2023). Canonical reactome pathways from MSigDB v2023.1.Hs release downloaded from gsea-msigdb.org website (Liberzon et al., 2011) were used to perform all enrichment analysis with enricher function of clusterProfiler R package (Wu et al., 2021). To simplify the enrichment results, enriched pathways with a *p* value ≥ 0.05 were selected, then a corresponding semantic similarity matrix was calculated and clustered by the binary cut algorithm with the R package simplifyEnrichment (Gu and Hübschmann, 2023). Three pathways with the lowest p-value in each group were retained to draw dotplots with the R clusterProfiler package (Wu et al., 2021).

### Statistical analysis

Results were represented as the mean +/- standard error of the mean (SEM). Nonparametric Mann-Whitney tests using Prism GraphPad software (San Diego, CA) were used to compare differences between variables.

## RESULTS

To deregulate Ig production in myeloma cells, we used a steric blocker ASO targeting a consensus RNA sequence surrounding the secreted “AAUAA” PAS in all *IGHG* subclasses (IgG-ASO) (Fig. S1). For passive diffusion through the plasma membrane, both IgG-ASO and an irrelevant control oligo (CTR) were synthetized as Vivo-Morpholino oligomers (Gene Tools, LLC). IL-6-independent (LP1) and IL-6- dependent (XG2 and XG6) myeloma cell lines were treated with ASO concentrations ranging from 1 to 6 µM, as indicated (Fig. 1). First, the impact of IgG-ASO treatment was assessed by quantifying secreted *IGHG* mRNA amounts after 48h treatment with low and high ASO doses close to the IC50 from each cell line. A significant and dose-dependent decrease in *IGHG* mRNAs was observed in all cell lines treated with IgG-ASO, as compared to control conditions (Fig. 1A-C). Consistent with decreased *IGHG* mRNA amounts, a drastic drop in IgG concentrations was found in culture supernatants (Fig. 1D-F). To a lesser extent, the targeting of secreted *IGHG* PAS sequence caused alternative polyadenylation with an increase in membrane *IGHG* mRNAs in IgG-ASO treated cells (Fig. 1G-I). Thus, efficient inhibition of IgG secretion was achieved upon treatment with an ASO targeting the secreted PAS of all IgG subclasses.

**Figure 1.**
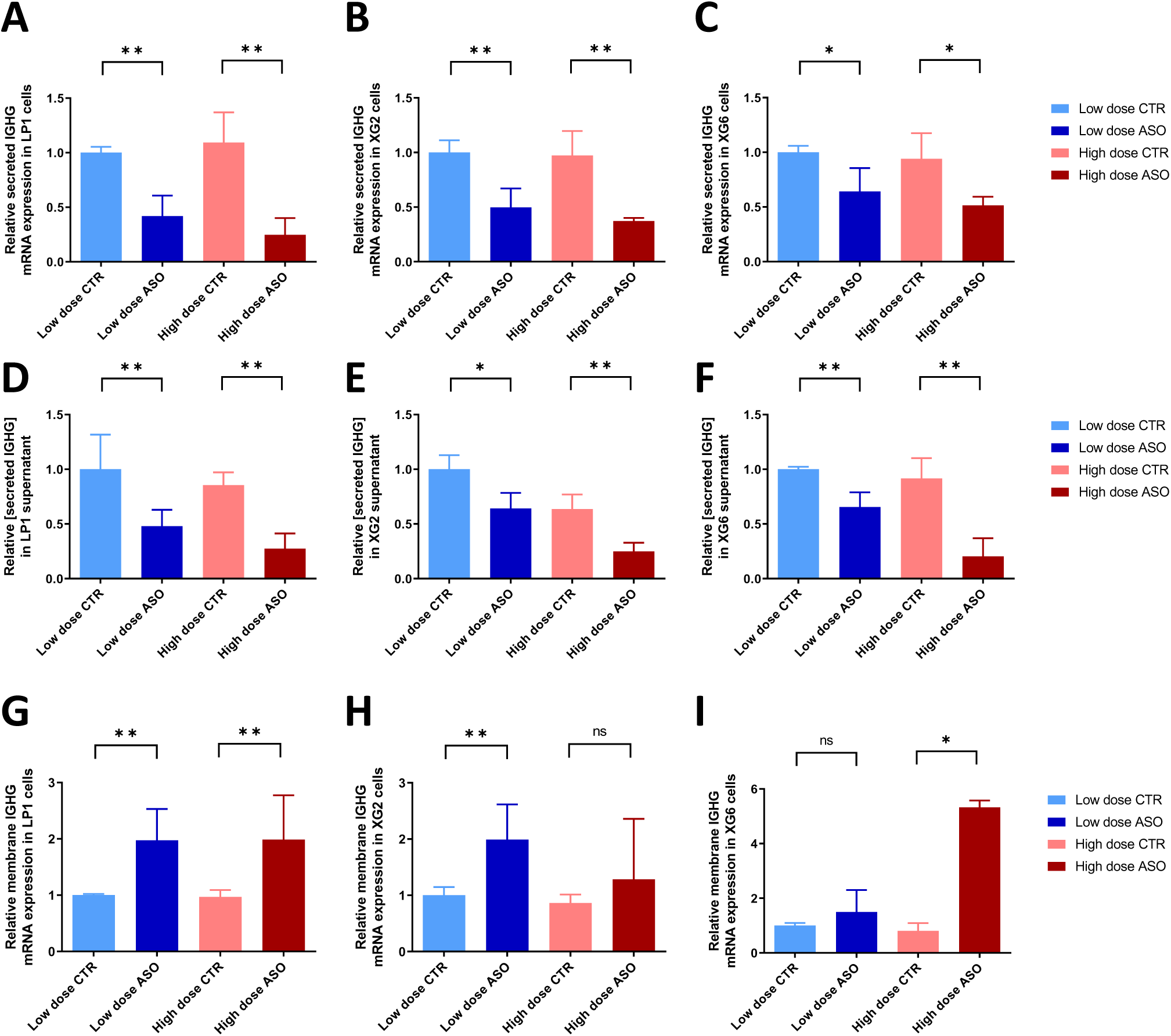
Decreased *IGHG* expression and IgG secretion upon treatment with an ASO targeting a consensus RNA sequence surrounding all secreted *IGHG* PAS. LP1 (A, D and G), XG2 (B, E and H) and XG6 (C, F and I) cells were with IgG-ASO (ASO) or with an irrelevant control ASO (CTR) for 48h. A-C) RT-qPCR analysis of the relative level of secreted *IGHG* mRNAs (sec-γHC), after normalization to the lowest dose of CTR. D-F) ELISA analysis of IgG secretion in culture supernatants. G-I) RT-qPCR analysis of membrane *IGHG* (mb-γHC) mRNA amounts after normalization to the lowest dose of CTR. For LP1 cells, low and high ASO doses correspond to 4 µM and 6 µM, respectively. For more ASO sensitive XG2 and XG6 cells, low and high ASO doses correspond to 1 µM and 3 µM, respectively. Data are from 2-4 distinct experiments, each performed with 2-3 culture replicates. *ns* (non significant), * (*p* value<0,05), ** (*p* value<0,01).

Next, we seeked to determine whether IgG-ASO treatment altered intracellular HC/LC stoichiometry at the protein level. Quantification was performed on cell pellets by Western Blot and ELISA (Fig. 2). Again, a strong decrease in IGHG protein levels was found in IgG-ASO treated cells compared to control conditions (Fig 2A-C). By contrast, the amount of IGL chains remained mostly similar in IgG-ASO- and CTR-treated cells (Fig. 2A, D-E). Then, Western Blots were performed under non-reducing conditions to investigate the extent of complete Ig, with proper HC and LC assembly, and free-LC species simultaneously (Fig. 3). As expected, lower amounts of complete Ig were found in IgG-ASO treated cells compared to control conditions (Fig. 3A upper bands). In addition, intracellular free-LC strongly accumulated as monomers and dimers in cells treated with IgG-ASO (Fig. 3A lower bands). Consistent with previous observations that normal and malignant PC produce excess of LC, these free-LC species were readily detectable in control conditions, allowing direct comparison of intracellular free-LC (monomers and dimers)/complete Ig ratios. We found a dose-dependent increase in free-LC/complete Ig ratios in cells treated with low (> 4.5-fold) and high (> 8.8-fold) IgG-ASO concentrations compared to controls (Fig. 3B). Free-Ig *lambda* light chains exhibited a propensity to be secreted mostly as dimers (Fig. S2) and we evaluated these amounts in culture supernatants. We found a nearly 2-fold increase in free-LC in cell culture supernatants form cells treated with IgG-ASO compared to control conditions (Fig. 3C). Thus, the reduction of IGHG proteins upon treatment with IgG-ASO markedly disturbed HC/LC stoichiometry while causing a strong intracellular excess a free-LC.

**Figure 2.**
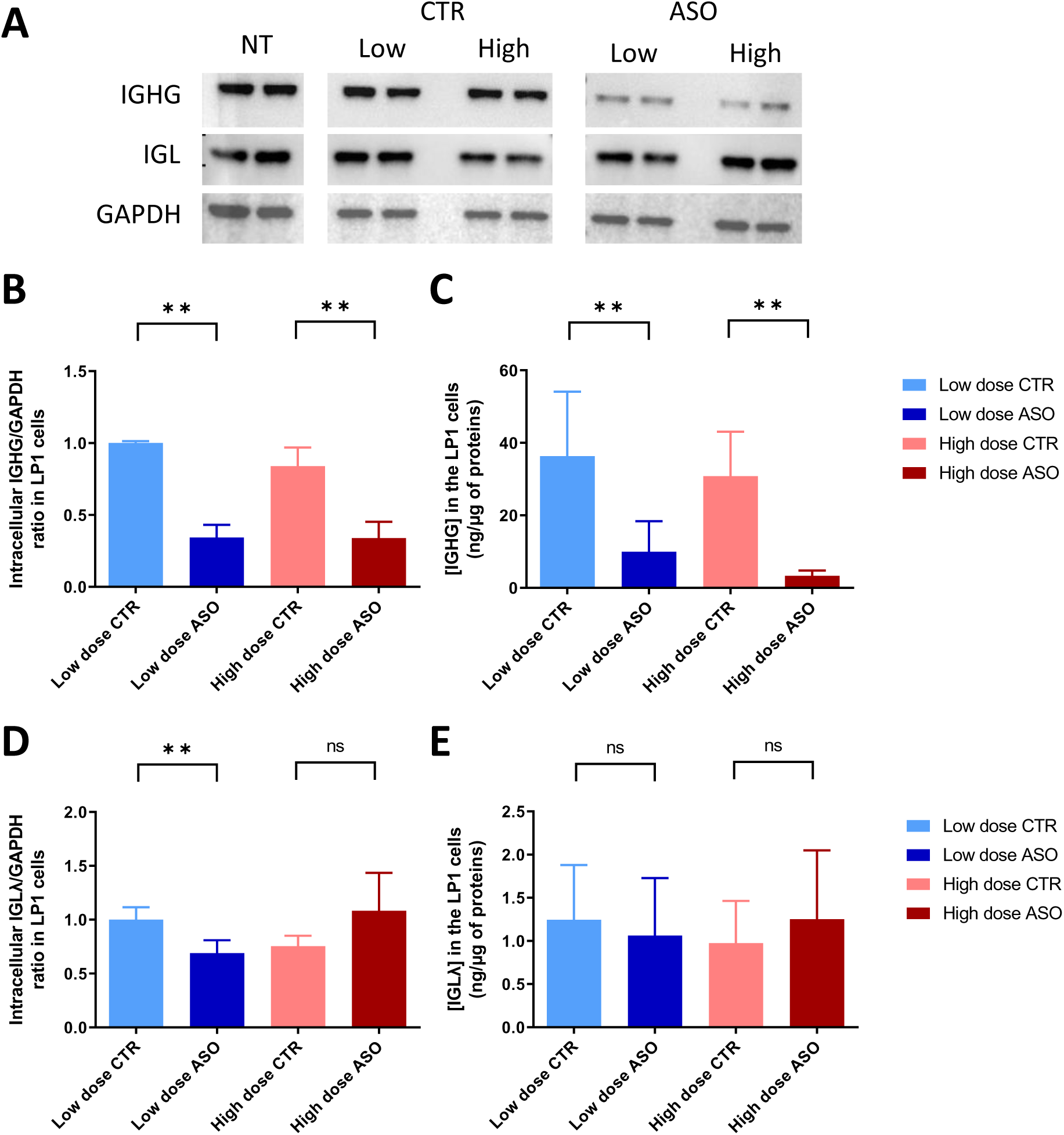
Specific decrease in IGHG protein levels without impairment of IGL production. LP1 cells were treated with IgG-ASO (ASO) or with an irrelevant control ASO (CTR) for 48h as in Fig. 1. A) Western Blot were performed on cell pellets with 10 µg of protein in each condition. A representative image is shown for the analysis of IGHG (up), IGL (middle) and GAPDH (down) expression. B and D) Quantification of IGHG (B) or (D) expression after normalization to GAPDH and to low dose CTR conditions. C and E) Quantification of IGHG (C) or IGL (E) expression on cell pellets after normalization to the amount of total proteins. Data are from 3 distinct experiments, each performed with 2 culture replicates. *ns* (non significant), ** (*p* value<0,01).

**Figure 3.**
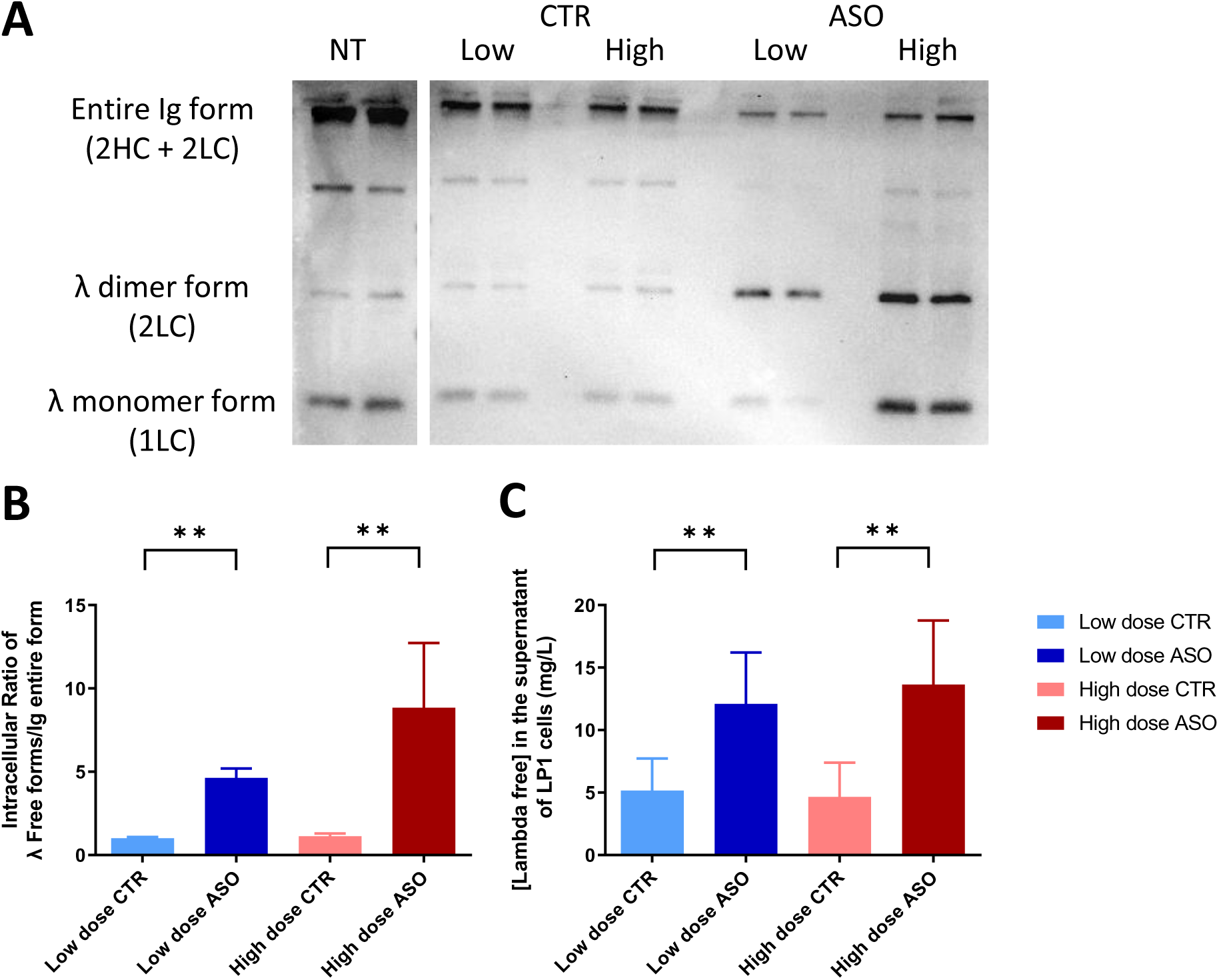
Treatment with IgG-ASO disturbed HC/LC ratios with a drastic increase in free-LC. LP1 cells were treated with IgG-ASO (ASO) or with an irrelevant control ASO (CTR) for 48h. A) Western Blot were performed in cell pellets (10 µg) under non-reduced conditions. Revelation was performed using anti-IGL antibody allowing simultaneous identification of entire Ig (2HC+2LC) or free-LC monomers (1LC) and dimers (2LC). A representative image is shown. B) Determination of free-LC/entire Ig ratios from band quantification of Western Blots. C) Quantification of free-LC in culture supernatants using the Optilite® Freelite® Lambda Free kit (The Binding Site™). Data are from 3 distinct experiments, each performed with 2 culture replicates. *ns* (non significant), ** (*p* value<0,01).

Then, RNA-seq analysis was performed to evaluate the global impact of disturbed HC/LC stoichiometry at the transcriptomic level. Poly(A)-enriched RNAs from IL-6-dependent XG6 cells treated with IgG-ASO and in control conditions were prepared for sequencing. There were 2,529 differentially expressed (DE) genes (adjusted *p* value < 0.01 and absolute log2foldchange > 0.5), with 882 upregulated and 1,647 downregulated in IgG-ASO treated cells compared to CTRL (Fig. 4A and Table S3). Of note, no major transcriptomic changes were observed by comparing CTR-treated cells to untreated cells, with a limited number of 244 DE genes (Fig. 4B and Table S4). Next, Reactome canonical pathway analysis was performed on a total number of 2393 DE genes exclusively included in IgG-ASO *vs* CTR conditions (Fig. 4B ; Table S5 and S6). Interestingly, several pathways involved in UPR and cell metabolism were found significantly altered in IgG-ASO treated cells compared to CTR cells (Fig. 4C; Table S6 and S7). As hallmarks of stressfull conditions particularly relevant for MM disease (Børset et al., 2022), the expression of genes such as *ATF4*, *HSPA5* and *HSP90B1* were significantly increased after treatment with IgG-ASO (Fig. 4A and Table S3). Thus, ASO-mediated disruption of HC and LC assembly provoked massive transcriptomic changes with altered expression of cell metabolism pathways, as well as genes involved in the maintenance of protein homeostasis.

**Figure 4.**
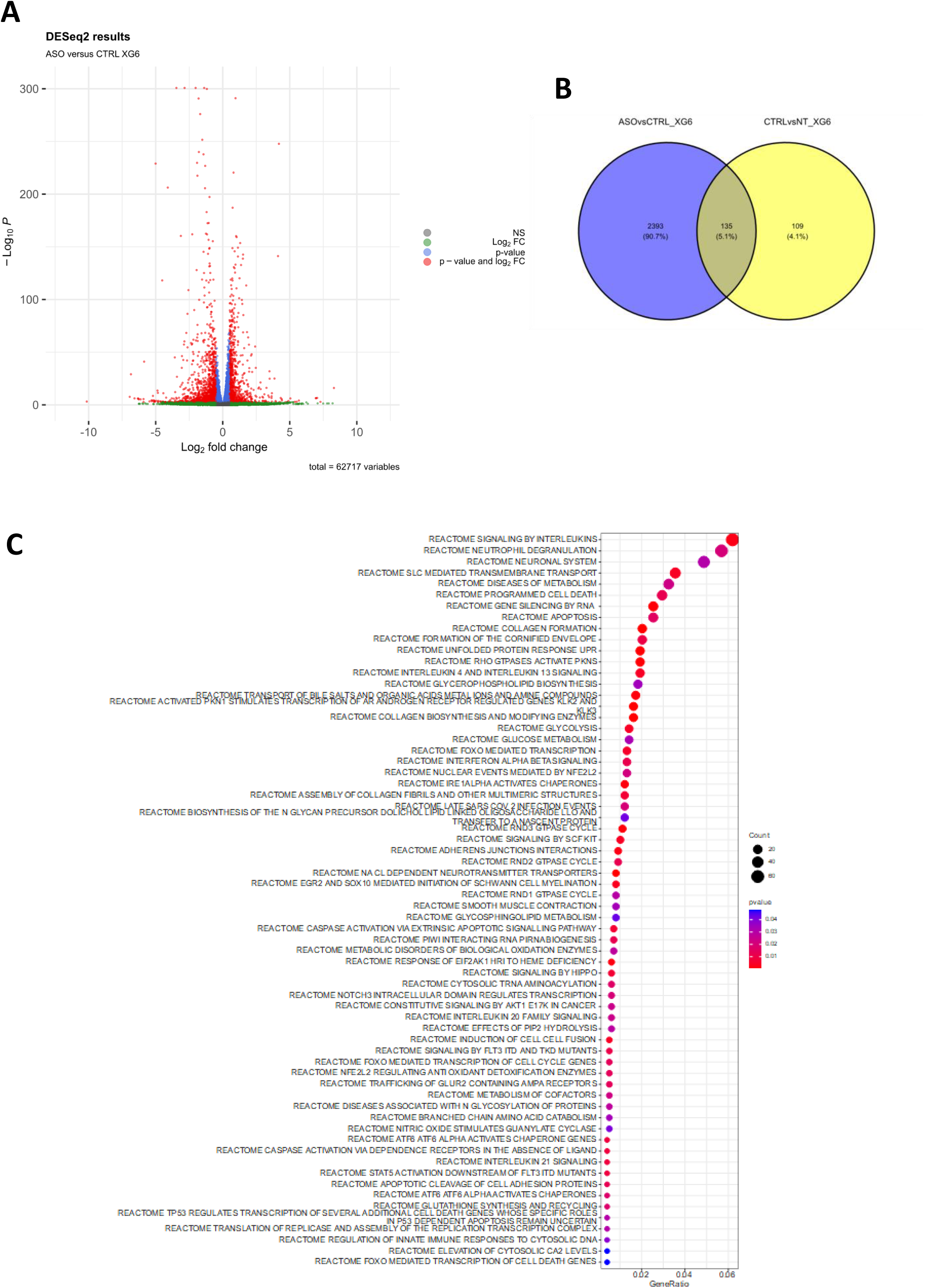
Major transcriptomic changes upon treatment with IgG-ASO with alteration of cell metabolism, UPR and cell death pathways. RNA-seq was performed on polyA-enriched RNA from XG6 cells treated or not with IgG-ASO or CTR (48h, 3µM ; n=3/condition). Paired-end (2x150bp) reads were processed through the bioinformatic pipeline nf-core/rnaseq v3.12 (Patel et al.) in a reverse strandedness mode. A) Volcano plot showing differentially expressed (DE) genes (adjusted *p* value < 0.01 and absolute log2foldchange > 0.5) as red dots, after comparison of IgG-ASO (ASO) and control (CTRL) treatments. B) Venn diagram showing that 2,293 DE were exclusively included in ASO *vs* CTRL conditions but not in CTRL *vs* NT (blue color). C) Canonical reactome pathways analysis from MSigDB v2023.1.Hs release showing enriched pathways with a *p* value ≤ 0.05 were selected. Three pathways with the lowest *p* value in each group were retained to draw dotplots with the R clusterProfiler package (Wu et al.).

Transcriptomic analysis also revealed that cell death and apoptotic pathways were strongly deregulated upon treatment with IgG-ASO (Fig. 4C; Table S3, S6 and S7). Indeed, viable cells were significantly reduced in XG6 and LP1 cells treated with IgG-ASO, compared to control conditions (Fig. 5A-C). Next, we determined the impact of IgG-ASO treatment on IgG-expressing primary cells isolated from bone marrow aspirates of MM patients (Table 1). Interestingly, lower frequency of CD38+ CD138+ population containing tumor PCs was achieved in all IgG-expressing MM patient cells (Fig. 5D). By contrast, we found no major effect in non-PC populations (Fig. 5E). Again, the treatment with IgG-ASO provoked a strong decrease in IGHG protein levels (Fig. 5F), with a dose-dependent inhibition in IgG secretion in culture supernatants over a 7-day period (Fig. S3). Thus, IgG-expressing primary MM cells exhibited high sensitivity to treatment with IgG-ASO whereas healthy cells were nearly unaffected.

**Figure 5.**
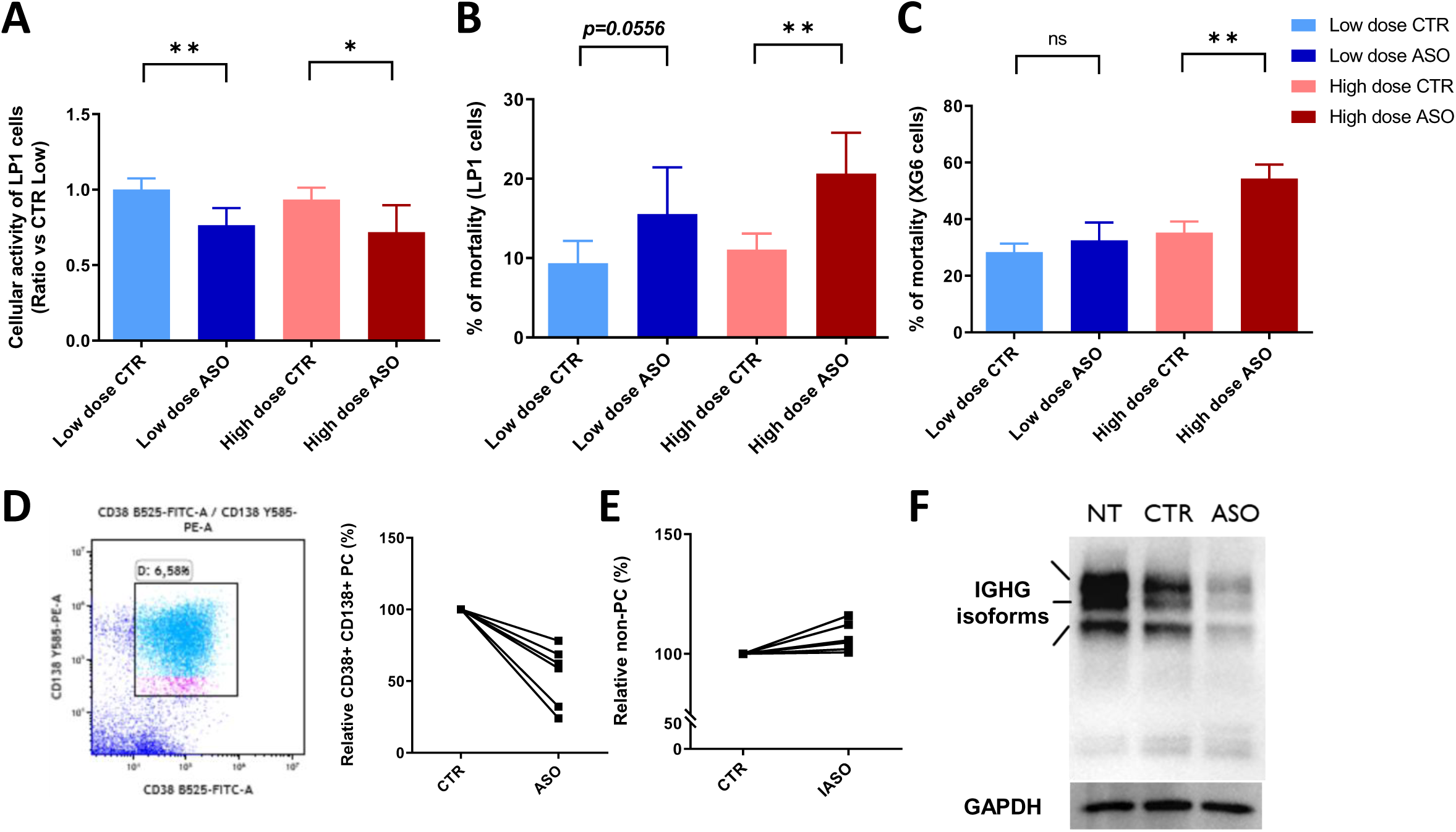
Impaired myeloma cell survival upon treatment with IgG-ASO. LP1 (A-B) cells were treated with low (4 µM) and high (6 µM) doses of IgG-ASO (ASO) and control (CTR) during 48h. A) MTS assays were performed during the last 3 hours of culture, and expressed after normalization to the lowest dose of CTR. B) Percentage of cell mortality obtained by counting cells after Trypan blue coloration on a Malassez slide. C) XG6 cells were treated with low (1 µM) and high (3 µM) doses of IgG-ASO (ASO) and control (CTR) during 48h and percentage of cell mortality was determined as in B. D) Representative dot plot obtained after 48h CTR treatment of primary MM patient cells isolated from bone marrow aspirates. The gating strategy for CD138+ CD38+ PCs is shown (left). The frequency of CD138+ CD38+ PCs was expressed for each MM patient sample treated with IgG-ASO (2.5 µM) after normalization to CTR (n=6, right). E) The frequency of non-PC populations (CD138-CD38-, CD138+ CD38- and CD138+ CD38-) was expressed for each MM patient sample treated as in D. F) A representative Western Blot image showing the strong decrease in IGHG isoforms in IgG- ASO treated MM patient cells. *ns* (non significant), * (*p* value<0,05), ** (*p* value<0,01).

**Table 1.** IgG-expressing primary myeloma cells isolated from bone marrow aspirates of MM patients.

| Name Id | Disease | Diagnosis/<br>Relapse | Subtype | Ig pic (g/L) | Infiltration (%) |
| --- | --- | --- | --- | --- | --- |
| DESMI | MM | Diagnosis | IgGκ | 22 | 65 |
| GERMI* | MM | Diagnosis | IgGλ | 51 | 49 |
| MARFA* | MM | Diagnosis | IgGλ | 30 | 37 |
| BOUCH* | MM | Relapse | IgGλ | 4 | 42 |
| BOUCL | MM | Diagnosis | IgGλ | 40 | 28 |
| PA1471* | MM | Diagnosis | IgGκ | / | 38 |
| PA1493* | MM | Diagnosis | IgGκ | / | 40 |
| PA1530* | MM | Diagnosis | IgGκ | / | 52 |

*\*MM patient cells used for flow cytometry analysis*

## DISCUSSION

RNA-based strategies are currently expanding worldwide but are still emerging in oncology. Unlike CRISPR/Cas technology, antisense strategies result in transient modifications of gene expression and lack mutagenic effects at the DNA level. In this study we describe the use of an ASO targeting a consensus RNA sequence found in all PAS from secreted *IGHG* subclasses. The aim of this antisense approach is to reduce IgG secretion, while sparing the production of other Ig isotypes (IgM, IgA, IgE). The other major interest lies in disrupting the assembly of Ig heavy and light chains in order to induce protein stress in PCs. Using myeloma cells as a monoclonal model of choice, we validated the efficacy of our strategy by showing a clear reduction in the quantity of secreted IgG. Interestingly, we show that a decrease in HC expression leads to an intracellular accumulation of free-LC, which disrupts cellular homeostasis and impairs myeloma cell survival.

The survival of plasma cells is intimately linked to the maintenance of intracellular protein stress through activation of the UPR. These stressful conditions associated with massive Ig synthesis must be finely regulated to avoid prolonged stress leading to cell death by apoptosis (Cenci et al., 2011). In the pathophysiology of MM, numerous examples in the literature highlight the marked sensitivity of myeloma cells to disturbances in protein homeostasis (Mimura et al., 2012; Obeng et al., 2006; Cenci et al., 2011). Previous work by Comenzo’s team, aimed at reducing the quantity of amyloidogenic LCs by administration of siRNA, also indicates toxicity following protein stress (Zhou et al., 2014; Ma et al., 2016). In these PC expressing entire Ig (with proper assembly of HC and LC), an exacerbated ER stress response was correlated with the accumulation of unpaired HC in the ER. Unlike our study showing a strong impairment of PC survival upon ASO-mediated inhibition of HC expression, these authors did not observe any toxicity when HC production was inhibited by a similar siRNA approach (Zhou et al., 2014). This discrepancy is not yet understood and may simply be due to a shorter incubation period with siRNA (24 h) than with ASO (48 h). Variability in HC/LC ratios between the different PC lines cannot be ruled out either, and this deserves further investigation.

In our study, the inhibition of secreted *IGHG* expression by ASO treatment resulted in a large accumulation of intracellular free-LC. Interestingly, the major transcriptomic changes observed indicated altered expression of genes involved in UPR, cell metabolism and apoptotic pathways. Although unpaired LCs are generally considered to be freely secreted (Katzmann et al., 2002; Zhou et al., 2014), our data suggest that an excess of free-LC can strongly disrupt protein homeostasis and alter the survival of PC-expressing entire Ig.

The main advantage of our strategy is that it can specifically inhibit the secretion of a particular HC isotype while largely sparing humoral immunity. We have previously demonstrated the value of this approach for inhibiting IgE secretion in allergy (Marchalot et al., 2021). Approaches specifically targeting the secretion of an HC isotype are also of great interest in clinical manifestations due to polyclonal Ig such as autoimmune diseases, IgA nephropathy or acute graft rejection. Regarding MGCS, the rationale for such an ASO-mediated strategy is particularly relevant for HC deposition diseases (HCDD) (Bridoux et al., 2017). However, care must be taken with regard to its therapeutic application in the treatment of MM or MGCS. Indeed, an excess of light chains in the circulation could damage organs, particularly with a risk of developing tubular kidney deposits as LC cast nephropathy (Hutchison et al., 2011). In addition, a free-LC escape can occur at relapse in MM patients (Dawson et al., 2007).

In conclusion, this study provides evidence for effective inhibition of IgG secretion using an ASO targeting the secreted PAS on *IGHG* transcripts. In addition, impaired survival of IgG-expressing myeloma cells correlated with an accumulation of free-LC. Altogether, ASO-mediated knockdown of HC expression could open new avenues for selective therapies, targeting PCs in a secreted Ig isotype- dependent manner.

## ACKNOWLEDGMENTS

We thank Anthony Vigier and Claire Carrion (CNRS UMR 7276 - INSERM U 1262 - Limoges University, Limoges, France) for their help with ELISA assays and flow cytometry experiments, respectively.

## Notes

**FUNDING** This work was supported by grants from INCa (PLBIO2022-110 to LD), ANR (2017-CE15-0024-01 to LD), Ligue Contre le Cancer (comités Corrèze, Haute-Vienne) and Fondation Française pour la Recherche contre le Myélome et les Gammapathies monoclonales (FFRMG). CH was supported by Région Nouvelle-Aquitaine and Limoges University.

### Competing Interest Statement

The authors have declared no competing interest.

