## Supplementary material for "Disruption of immunoglobulin heavy and light chain assembly by antisense oligonucleotides impairs protein homeostasis and myeloma cell survival": Horiot et al sup.docx

**SUPPLEMENTAL INFO**

Table S1. List of antibodies used in this study

| Antibody used | Coupled | Reference | Assay | Dilution |
| --- | --- | --- | --- | --- |
| Mouse anti-human CD138 Ab | PC5 | Beckman Coulter IOTest, Ref#A54191 | Flow cytometry | 1/10 |
| Mouse anti-human CD38 Ab | BV421 | Biolegend #356617 | Flow cytometry | 1/10 |
| Goat anti-human IgG | HRP | Southern Biotech #2040-05 | WB | 1/2000 |
| Polyclonal Rabbit anti-human Lambda light chains | / | Dako #A0194 | WB | 1/2000 |
| Goat anti-Rabbit IgG (H+L) Mouse/Human ads-HRP | HRP | Southern Biotech #4050-05 | WB | 1/4000 |
| Mouse anti-β-actin Ab | / | Sigma #A5441 | WB | 1/10000 |
| Goat anti-Mouse IgG | HRP | Cell Signaling #7076S | WB | 1/2000 |
| GAPDH (D16H11) XP® Rabbit mAb | HRP | Cell Signaling #8884S | WB | 1/1000 |
| Goat anti human IgG Ab (coating) | UNLB | Southern Biotech #2040-01 | ELISA | 1/1000 |
| Goat anti human IgG Ab (detection) | AP | Southern Biotech #2040-04 | ELISA | 1/1000 |
| Goat anti human lambda Ab (coating) | UNLB | Southern Biotech #2070-01 | ELISA | 1/1000 |
| Goat anti human lambda Ab (detection) | AP | Southern Biotech #2070-04 | ELISA | 1/1000 |

Table S2: List of primers used for qPCR

| Gene | Forward primer | Reverse primer |
| --- | --- | --- |
| secreted *IGHG1* | 5'-AAGCTCACCGTGGACAAGAG-3' | 5'-CGGCCGTGGCACTCATTTA-3' |
| membrane *IGHG1* | 5'-AAGCTCACCGTGGACAAGAG-3' | 5'-AGCACACGCTTAACAGGAAGA-3' |
| *IGL* | 5'-ATGGCCTGGDYYVYDCTVYTYCT-3' | 5'-CTCCCGGGTAGAAGTCACT-3' |

*with D = A or G or T, Y = C or T, and V = A or C or G.*

**Figure S1. Alignment of PAS from all secreted *IGH* subclasses**

A) Alignment of *IGHG* DNA sequences encompassing the secreted *IGHG* PAS (blue rectangle). Blue letter indicates nucleotide changes in sequence samples. The IgG-ASO sequence that binds to a conserved RNA sequence found in all IGHG subclasses is shown. B) Alignment of secreted PAS from all IGH subclasses showing the isotype specificity of IgG-PAS compound.

**Figure S2. Quantification of entire Ig and free-LC in culture supernatants**

LP1 cells were treated with IgG-ASO (ASO) or with an irrelevant control ASO (CTR) for 48h. Western Blot were performed in cell culture supernatants pellets under non-reduced conditions. Revelation was performed using anti-IGL antibody allowing simultaneous identification of entire Ig or free-LC monomers and dimers as in Fig. 3A. A representative image out of 2 performed is shown.

**Figure S3. Time course analysis of IgG secretion in culture supernatants from MM patient cells treated with ASO.**

Cell cultures were done over a 7 days with a low (2.5µM) or high (5 µM) dose of CTR or IgG-ASO molecules**.** ELISA analysis of IgG secretion (µg/mL) in culture supernatants from GERMI (A) and MARFA (B) patients. Data are represented as cell culture duplicates.
