## Supplementary material for "Disruption of immunoglobulin heavy and light chain assembly by antisense oligonucleotides impairs protein homeostasis and myeloma cell survival": Horiot et al sup.pptx

### Slide 1
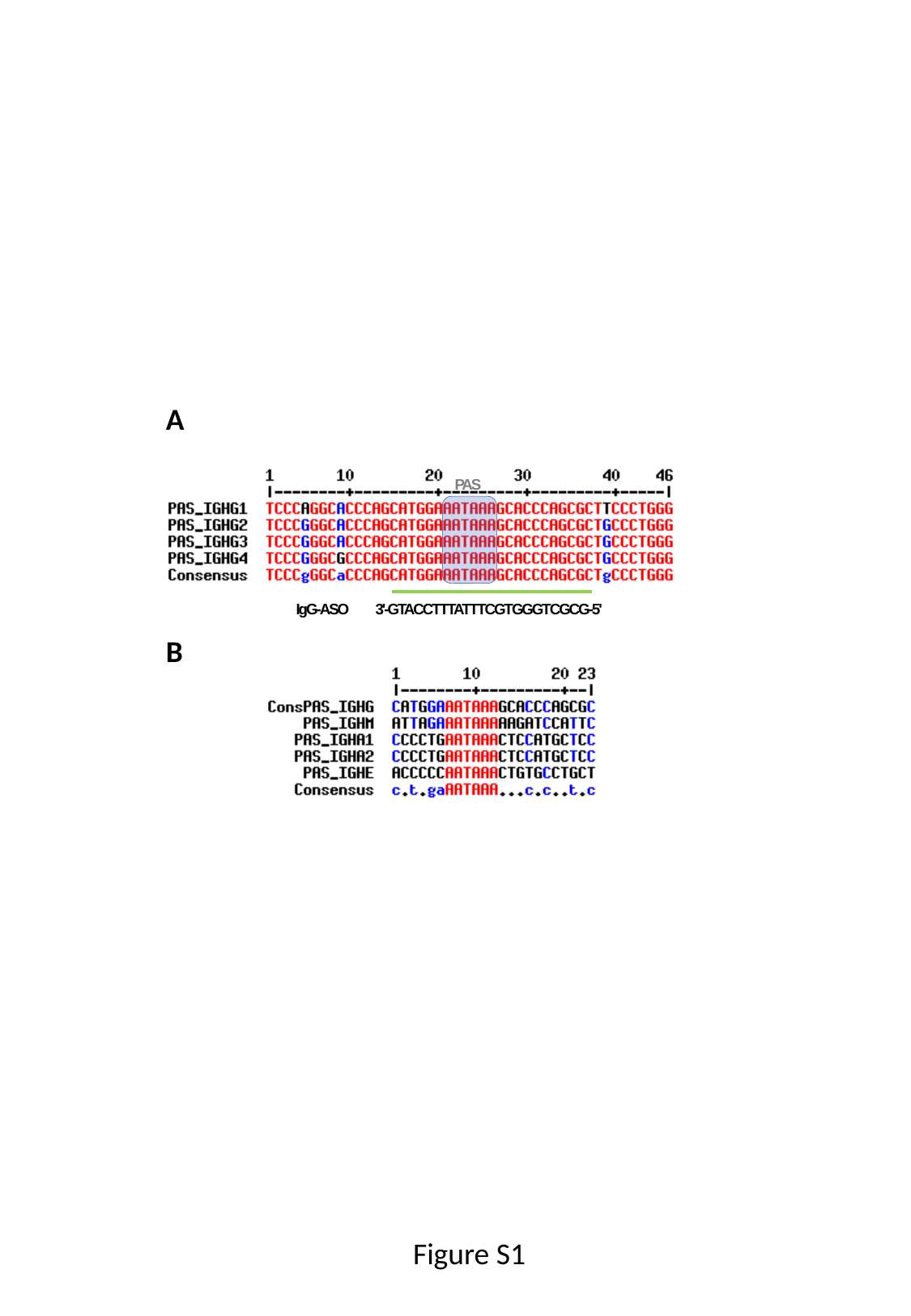

A
PAS
3'-GTACCTTTATTTCGTGGGTCGCG-5'
IgG-ASO
B
Figure S1

### Slide 2
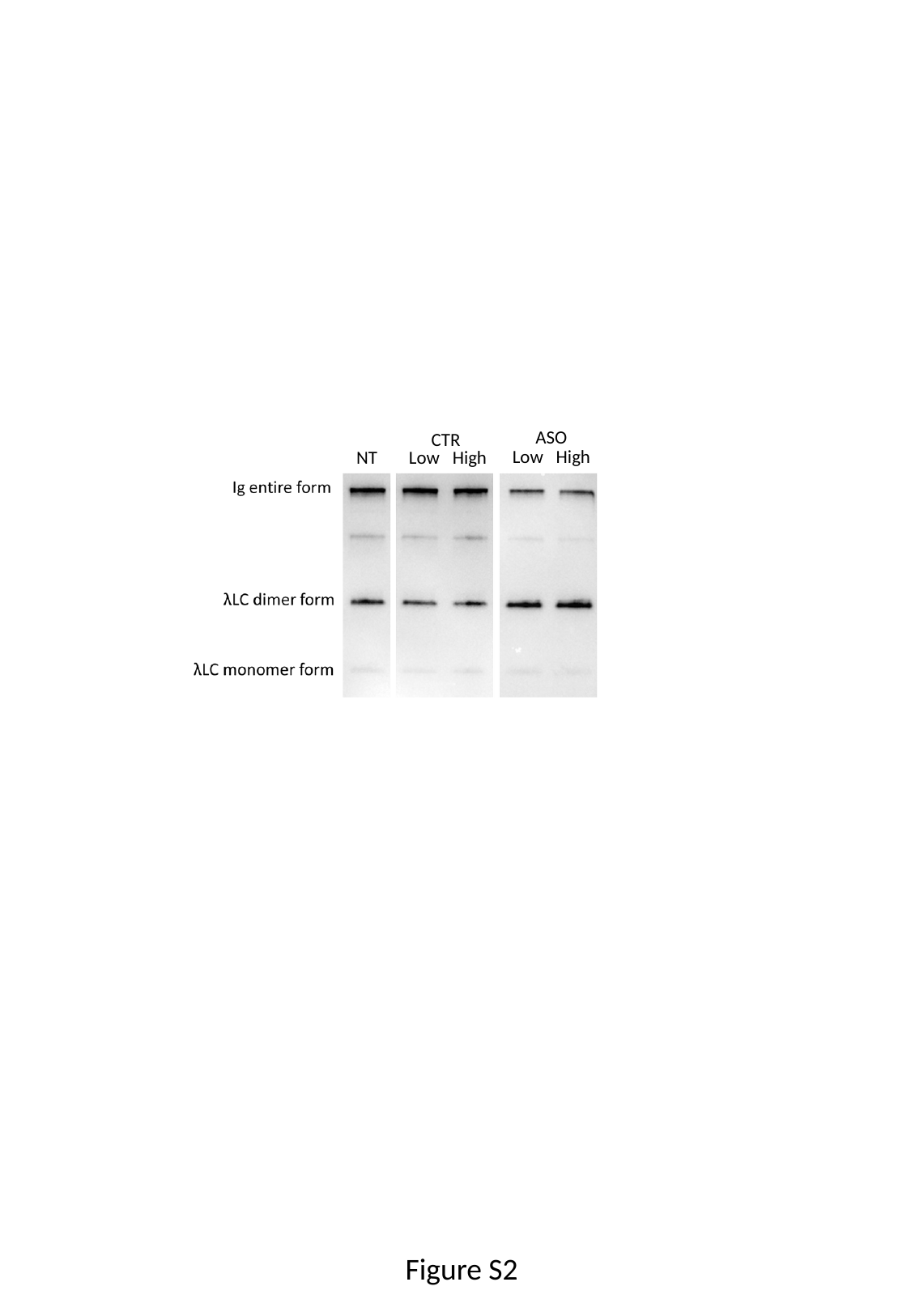

ASO
CTR
Low High
NT
Low High
Figure S2

### Slide 3
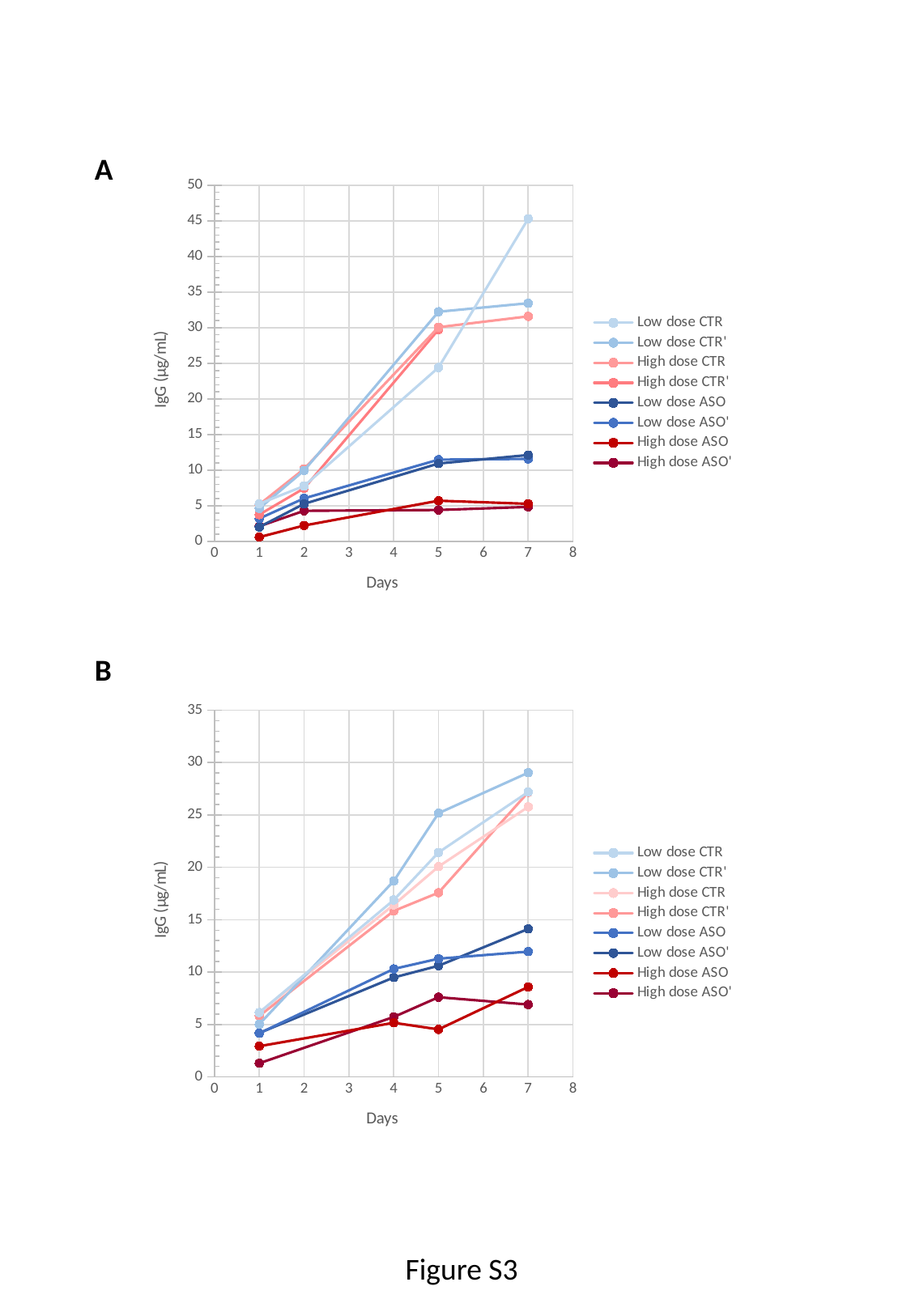

A
#### Chart
| Category | | | | | | | | |
|---|---|---|---|---|---|---|---|---|B
#### Chart
| Category | | | | | | | | |
|---|---|---|---|---|---|---|---|---|Figure S3
